# Experience-specific anterograde amnesia: memory reacquisition deficit phenomenon and its characterization in the vertebrate learning model

**DOI:** 10.1101/461616

**Authors:** A. A. Tiunova, D. V. Bezryadnov, D.R. Gaeva, V.S. Solodovnikov, K.V. Anokhin

**Affiliations:** P. K. Anokhin Research Institute of Normal Physiology, 125315 Moscow, Russia; National Research Center “Kurchatov Institute”, 123182 Moscow, Russia; Lomonosov Moscow State University, 119991 Moscow, Russia

**Keywords:** memory, amnesia, NMDA receptors, protein synthesis

## Abstract

A common assumption from experiments that interfere with memory consolidation is that the resultant amnesia returns the brain of an animal to a tabula rasa state with respect to disturbed experience. However, recent studies in terrestrial snail classical conditioning revealed an odd phenomenon: animals were unable to relearn conditioned avoidance of specific food after this memory had been impaired by protein-synthesis inhibitors or N-methyl-D-aspartate (NMDA) receptor antagonists. Here we examined whether such specific memory reacquisition deficit can also be observed in vertebrate learning. We trained day-old chicks in a one-trial passive avoidance task by presenting them a bead of a specific color covered with a repellent substance, methyl anthranilate. Training was preceded by administration of the protein synthesis inhibitor anisomycin or the NMDA receptor antagonist MK-801. Both drugs produced permanent amnesia, and no spontaneous recovery of memory was observed. A second training was given to the amnestic animals either using a bead of the same color (retraining) or a new color (novel training). The interval between the first and second training was 2 or 24 h, and the retention test was given from 30 min to 48 h after the second training. Retraining of the amnestic chicks with the bead that was presented during the initial training failed to produce new avoidance memory for this stimulus at all the between-training and training-to-test intervals. This memory reacquisition deficit was specific and was not transferred to a new conditioned stimulus, which was readily learned. We suggest that such pharmacologically induced experience-specific anterograde amnesia might reflect general properties of normal memory allocation, and we discuss its possible neural bases.

## Introduction

The neurobiology of memory has gained considerably from methods of experimental interference with memory consolidation or reconsolidation (Schafe et al. 2001; Lee et al. 2004; Debiec et al. 2002; Ben Mamou et al. 2006; Crowe et al. 2008; Solomonia and McCabe 2015). Though it is still disputable whether the interference-induced deficit reflects memory loss or a retrieval failure, it is presently well established that such treatments impair performance on subsequent retention tests, thus producing amnesia (Hardt et al. 2009; Nader 2015; Davis and Zhong 2017). Until recently, a natural assumption from these experiments has been that amnesia returns the brain to a sort of a tabula rasa state with respect to the impaired experience, making the animal ready for the subsequent reacquisition of the lost memory.

However, recent data obtained in terrestrial snails demonstrated a peculiar phenomenon of irreversibility of amnesia produced by protein-synthesis inhibition during memory consolidation or reconsolidation (Solntseva et al. 2007; Solntseva and Nikitin 2009). The snails trained in a food-aversion model have been made amnestic by a protein-synthesis inhibitor and were later trained again for the same conditioned stimulus (CS). Surprisingly, 3 consecutive days of the repeated training failed to produce a new aversion behavior. At the same time, the snails were able to acquire a new aversion memory for a different CS (Solntseva et al. 2007; Solntseva and Nikitin 2009). The same phenomenon was also observed in amnesia produced by protein synthesis inhibitors or N-methyl-D-aspartate (NMDA) receptor antagonists administered during memory reconsolidation (Nikitin and Solntseva 2013; Nikitin et al. 2016; Nikitin et al. 2017).

In the present study we tested whether such memory reacquisition deficit can be also observed in the vertebrate learning model. For this purpose, we used a one-trial passive avoidance learning task in new-born chicks (*Gallus gallus domesticus)*. This learning model, which is based on the innate predisposition of young chicks to peck at small objects and to memorize their characteristics, has been widely used to study neural mechanisms of memory consolidation and reconsolidation (Ng et al. 1997; Rose 2000; Matsushima et al. 2003; Crowe et al. 2008). Using this model, it has been shown that administration of protein synthesis inhibitors or NMDA receptor antagonists at the time of training produces permanent amnesia (Burchuladze and Rose 1992; Rickard et al. 1994; Freeman et al. 1995; Anokhin et al. 2002). NMDA antagonists were shown to affect consolidation of long-term memory for passive avoidance in chicks if injected within an interval from 30 min before to 10 min after the training (Burchuladze and Rose 1992, Steele and Stewart 1993, Rickard et al. 1994). Intraventricular administration of protein synthesis inhibitor anisomycin between 30 min pre-training and 75 min post-training also prevented consolidation of long-term passive avoidance memory (Freeman et al. 1995). Injection of anisomycin 5 min prior to training produced amnesia starting from 60 min post-training, and the recall deficit was stable for at least 48 h (Anokhin et al. 2002). Here we examined whether such amnesia can be overcome by retraining of the amnestic animals.

## Results

### I. Anisomycin-induced amnesia

Protein synthesis inhibitor anisomycin (ANI) was administered intraventricularly (80 μg/chick, 5 μl/hem) 5 min before the first training. The second training given to the anisomycin-injected chicks was considered either as retraining (presentation of a bead of the same color as at the first training) or as novel training (presentation of a bead of a new color). During the second training most of the animals pecked at the presented bead (58%-83% for the bead of the same color and 61%-93% for the bead of a new color). There were no statistically significant differences between the proportions of chicks that pecked at the beads of the same or of a novel color. Thus, anisomycin impaired memory after the first experience, thereby allowing us to train the chicks for the second time using the same CS.

#### 1. Anisomycin-induced memory reacquisition deficit 24 hours after the initial training

In these reacquisition experiments, chicks received the second training 24 h after the initial training and were tested 0.5 h, 2 h, or 24 h later. Groups that received only initial training (Saline and ANI) were tested at 24.5, 26, or 48 h after the initial training, respectively. Administration of anisomycin during the initial training significantly impaired the avoidance behavior 24.5 h later (Fig. 1a, ANI group compared with the Saline group, *P*<0.05). Repeated training of anisomycin-injected animals with the same color bead failed to restore the aversion of this bead in the test 0.5 h later, their avoidance remaining significantly lower than in the saline group (Fig.1a, Retraining group, *P*<0.05). However, training of anisomycin-injected chicks with the bead of a new color produced clear avoidance of this novel stimulus 0.5 h later (Fig.1a, Novel group, *P*<0.001 compared with the Retraining group).

**Fig.1.**
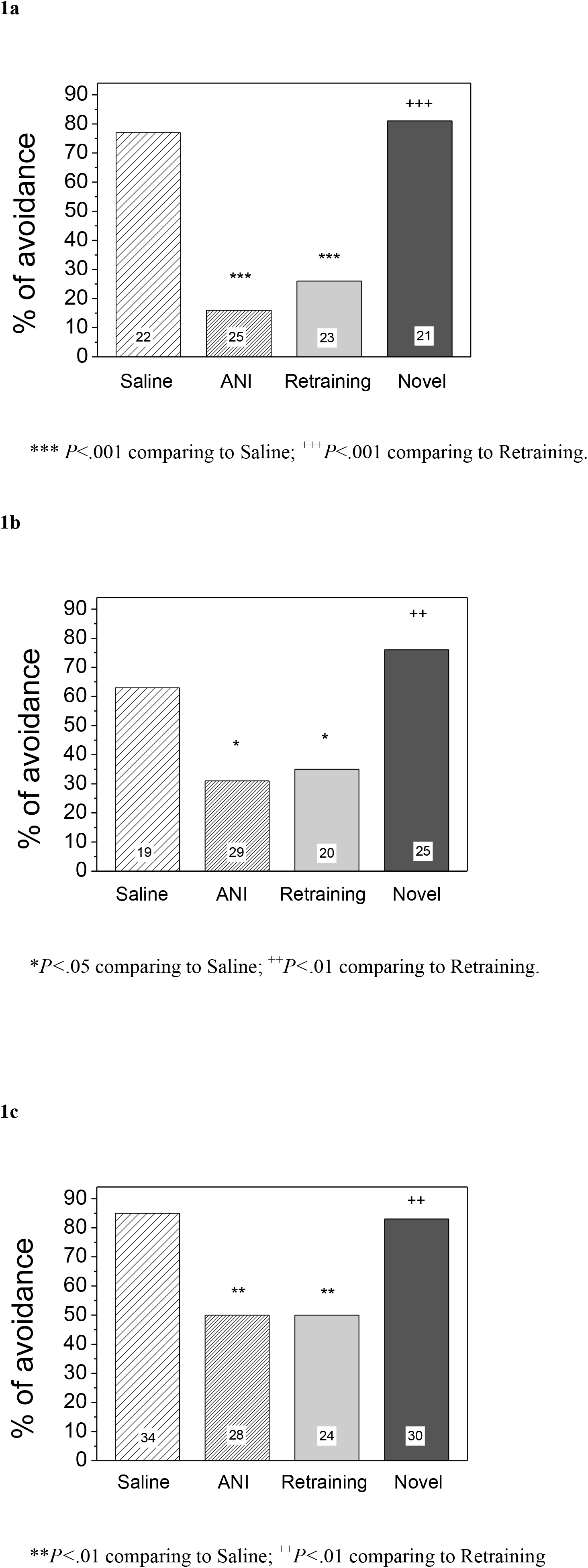
Avoidance level at the test given 0.5 h (**a**), 2 h (**b**) or 24 h (**c**) after the second training for the same stimulus (Retraining) or for a novel one (Novel). Initial training was coupled with anisomycin (ANI) administration; between-training interval 24h. Number of chicks is shown in the bars.

Administration of anisomycin during the initial training also impaired avoidance response in chicks in the 26 h test, ie, 2 h after the second training (*P*<0.05 compared with Saline). Repeated training of anisomycin-injected animals with the same stimulus failed to enhance their avoidance of the same CS 2 h later (Fig.1b, Retraining group, *P*<0.05 compared with Saline). Furthermore, the test given 24 h after the second training (48 h after the initial training) produced similar results (Fig. 1c). Thus, retraining of chicks 24 h after anisomycin-induced memory disruption failed to produce new short-term and long-term memory for the same CS, though training for a new CS led to formation of unambiguous memory.

#### 2. Anisomycin-induced memory reacquisition deficit 2 hours after the initial training

In the next experiment we examined whether a shorter between-training interval could result in effective reacquisition of the impaired memory. For this, the second training was conducted 2 h after the first one and the recall test was given 2, 24, or 48 h after the second training. In the test given 2 h after the second training, the avoidance level in the anisomycin group was significantly lower than in the saline group, showing that animals developed amnesia (*P*<0.01, Fig. 2a).

**Fig.2.**
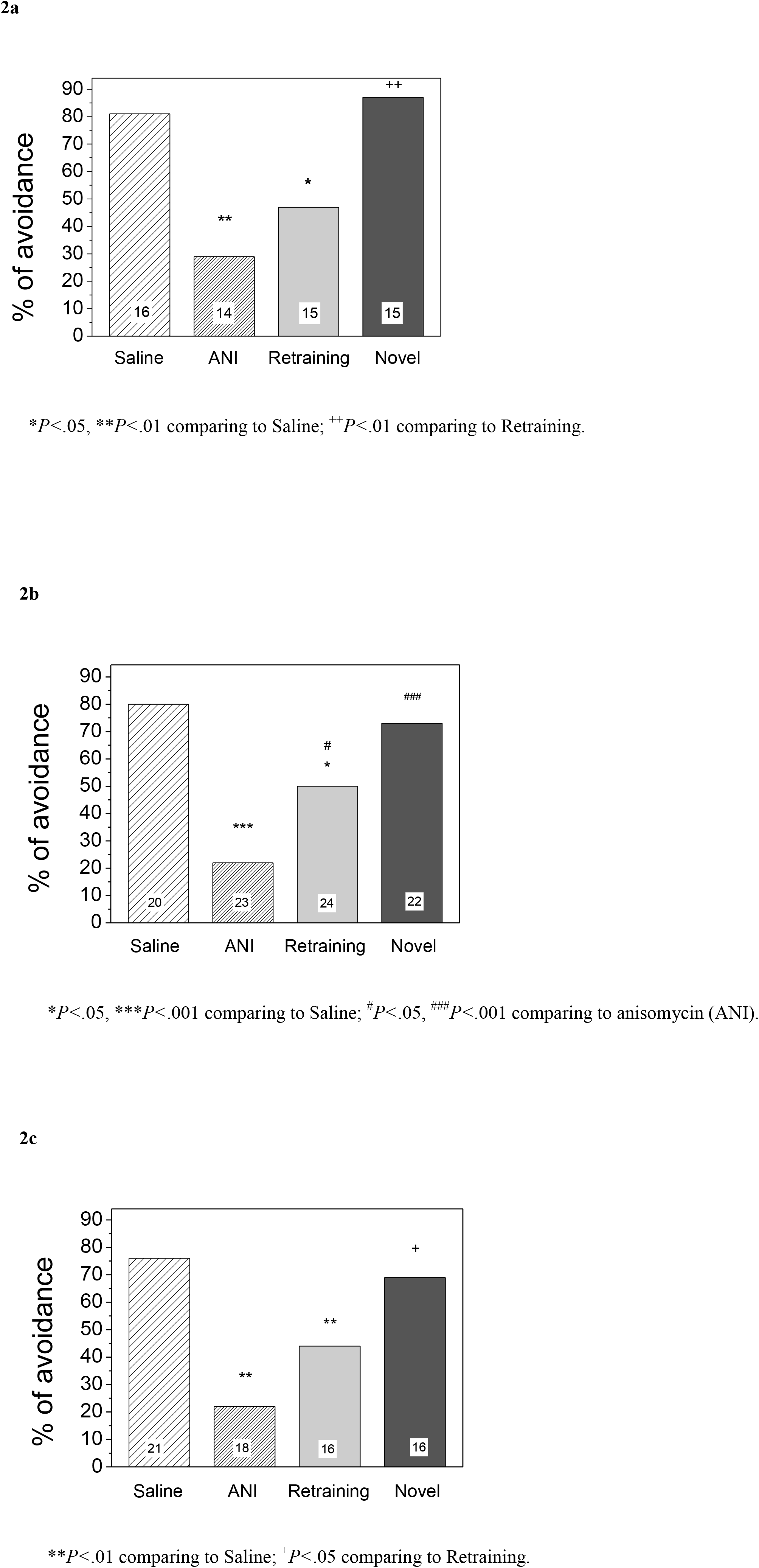
Avoidance level at the test given 2 h (**a**), 24 h (**b**) or 48 h (**c**) after the second training for the same stimulus (Retraining) or for a novel one (Novel). Initial training was coupled with anisomycin (ANI) administration; between-training interval 2h. Number of chicks is shown in the bars.

Repeated training of the anisomycin-injected chicks did not produce significant avoidance of the bead of the same color, whereas training them with a new stimulus resulted in as high avoidance as in the saline group (Fig. 2a, *P*<0.01 compared with the Retraining group). At the retention test both 24 h and 48 h after the second training, the avoidance remained high in the saline groups and was significantly impaired in the anisomycin groups (*P*<0.001, Fig.2b; *P*<0.01, Fig.2c). Again, training with a new stimulus produced normal memory that did not differ from the respective saline group and was higher than in the corresponding anisomycin group (*P*<0.001, Fig.2b; *P*<0.05, Fig.2c). In the chicks trained with the same bead (retraining groups) the avoidance level 24 h after the second training was statistically different from both the saline and anisomycin groups (*P*<0.05, Fig. 2b), whereas 48 h after the second training the avoidance level was significantly lower than in the saline group (*P*<0.01) and did not differ from the anisomycin group (Fig. 2c). Thus, repeated training failed to produce novel avoidance memory to the stimulus presented 2 h earlier in combination with the protein synthesis inhibitor anisomycin.

### II. MK-801–induced amnesia

NMDA receptor antagonist MK-801 (0.4 mg/kg) was injected intraperitoneally 5 min prior to the first training except in one experiment when it was administered intraventricularly (15 nmol 5 min pre-training). As in the anisomycin experiments, most of the MK-801–injected animals (51%-74%) demonstrated no avoidance reaction during the test, which allowed us to train them repeatedly with the same stimulus.

#### 1. MK-801–induced memory reacquisition deficit 24 hours after the initial training

In these experiments the second training was conducted 24 h after the initial training. The test was given 0.5 h, 2 h, or 24 h after the second training.

In the 0.5 h test, avoidance in the retraining group did not differ from the avoidance of chicks in the MK-801 group (Fig.3a) and was significantly lower than in the saline and Novel groups (*P*<0.01, Fig.3a). In the 2 h test, the avoidance in the retraining group also did not differ from that of the MK-801 group, and was significantly lower than in the saline group (*P*<0.01). Training of chicks with a novel stimulus produced significantly higher avoidance than retraining with the same stimulus (*P*<0.05, Fig.3b).

**Fig.3.**
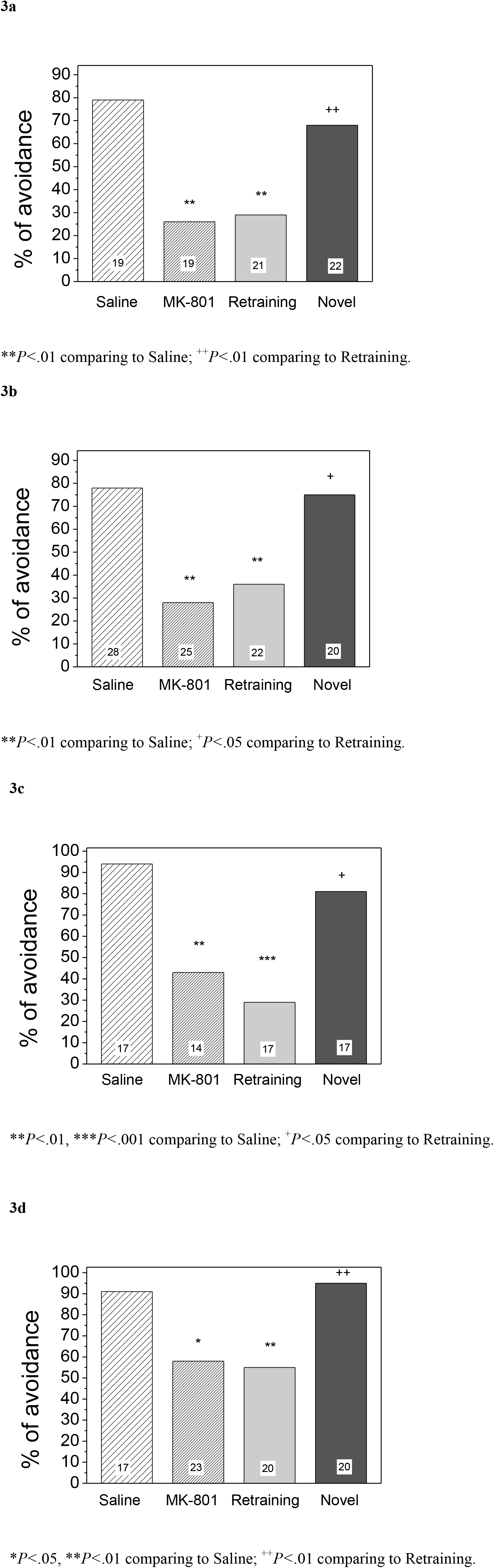
Avoidance level at the test given 0.5 h (**a**), 2 h (**b**) or 24 h (**c**, **d**) after the second training for the same stimulus (Retraining) or for a novel one (Novel). Initial training was coupled with intraperitoneal (**a, b, c**) or intraventricular (**d**) MK-801 administration; between-training interval 24h. Number of chicks is shown in the bars.

In the experiments where the test was given 24 h after the second training, we checked the effects of both central and systemic injection of MK-801. In both cases, repeated training failed to increase the avoidance level compared with the MK-801 group, while training with a novel stimulus produced a pronounced avoidance (*P*<0.05, Fig.3c; *P*<0.01, Fig.3d) that did not differ from the avoidance level in the respective saline group (Fig. 3c, d). Thus, 24 h after the training accompanied by NMDA receptor antagonist MK-801, repeated training with the same stimulus failed to produce short-term or long-term memory. At the same time, training with a novel stimulus resulted in the development of normal avoidance memory to this object.

#### 2. MK-801–induced memory reacquisition deficit 2 hours after the initial training

Next, we examined whether chicks can relearn the task at a shorter interval after the amnesia onset. A second training was carried out 2 h after the first training that was accompanied by MK-801. The tests were given 2 h or 24 h after the second training. In both tests the avoidance level in the retraining group was significantly lower than in the saline group (*P*<0.001, Fig.4a; *P*<0.05, Fig.4b). In contrast, chicks trained for a new stimulus demonstrated high avoidance levels (compared with the Retraining group: *P*<0.001, Fig.4a; *P*<0.05, Fig.4b).

**Fig.4.**
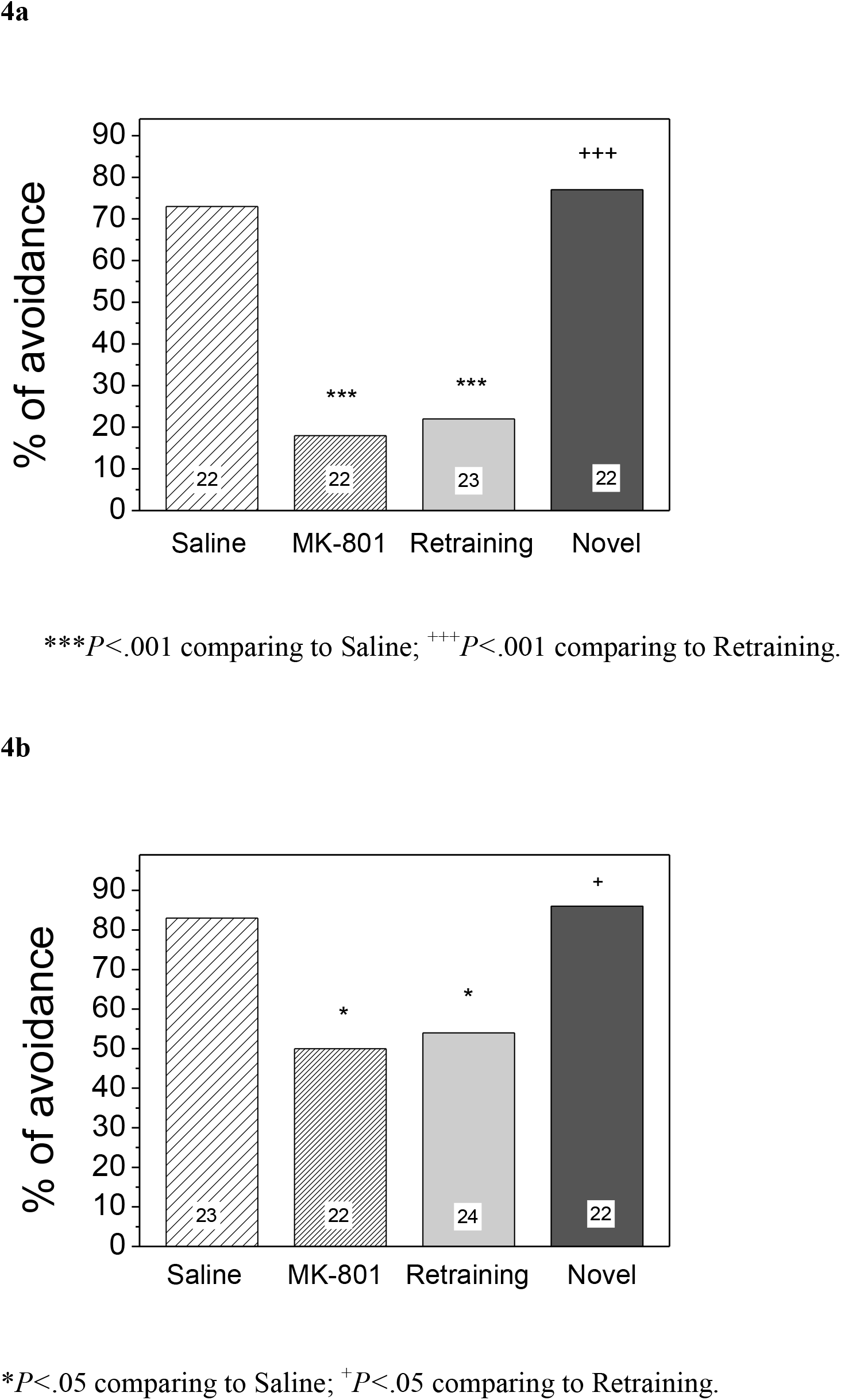
Avoidance level at the test given 2h (**a**) or 24h (**b**) after the second training for the same stimulus (Retraining) or for a novel one (Novel). Initial training was coupled with MK-801 administration; between-training interval 2h. Number of chicks is shown in the bars.

## Discussion

Our aim in the present study was to examine reversibility of experimental amnesia to a specific CS produced in chicks by pharmacological interference with memory consolidation. Young chicks were injected with a protein synthesis inhibitor, anisomycin, or an NMDA receptor antagonist, MK-801, prior to being trained to avoid a conspicuous color bead covered with an aversive substance. This treatment produced amnesia, which is in line with previous data (Burchuladze and Rose 1992; Rickard et al. 1997). In particular, amnesia as a result of such treatment was permanent and no spontaneous recovery of memory was observed. Due to this stable amnesia we were able to train the amnestic chicks again using the same CS to check whether they are capable of relearning anew the forgotten experience.

Altogether, the second training failed to produce a new avoidance memory for the same CS in the amnestic chicks at all between-training and training-to-test intervals.

This effect could not be attributed to a sustained influence of drugs during the second training. First, the minimal interval between drug administration and the repeated training was 2 h, which is longer than the known time window for the action of both anisomycin and MK-801 (Burchuladze and Rose 1992; Patterson et al. 1986). Second, at the same time the animals with impaired memory were able to learn and recall normally a novel stimulus that differed from the first one only by its color (Novel groups). Comparison of avoidance scores in the retraining and Novel groups implies selective failure of amnestic animals to reacquire memory only for the specific object used during the initial learning.

In these experiments we used 2 drugs that influence different stages of memory formation. The non-competitive NMDA receptor antagonist MK-801 is effective prior to or shortly after training, and affects early synaptic stages of memory formation (Burchuladze and Rose 1992; Rogers 1993; Rose 2000). Protein-synthesis inhibitor anisomycin impairs translation and does not affect short-term memory (Mark and Watts, 1971; Patterson et al. 1986). Here, we demonstrate that both drugs produce amnesia that is not reversed by the second training, either 2 h or 24 h after the amnesia onset.

A similar specific memory reacquisition deficit was reported previously in mollusks. Interference with consolidation or reconsolidation of aversive memory in the terrestrial snail (*Helix lucorum*) caused not only impairment of conditioned food-aversion memory but also the inability to relearn the avoidance of the specific stimulus that has been used during the initial conditioning (Solntseva and Nikitin 2010, 2012). This specific defect, called by the authors “irreversibility of amnesia,” was produced by antagonists of NMDA receptors or inhibitors of protein synthesis both during consolidation and reconsolidation of memory (Nikitin and Solntseva 2013; Nikitin et al. 2016; Nikitin et al. 2017). Intriguingly, the reversibility of amnesia (ie, the capability of amnestic animals to reacquire the lost memory) depended on the interval between the onset of amnesia and repeated training. If the interval between the onset of amnesia and the second training was less than 10 days, snails were able to memorize anew the same stimulus. However, if the interval was more than 10 days, the second training to the same stimulus did not produce new memory (Solntseva et al. 2007; Solntseva and Nikitin 2010). These data imply that in this model pharmacologically induced amnesia is not merely an absence of memory but rather a developing process that gradually unfolds to become irreversible. However, in our experiments we did not find strong evidence for such dynamic unfolding of amnesia. Memory reacquisition deficit was expressed in chicks as early as 2 h after impairment of the initial memory.

Neurobiological mechanisms underlying memory reacquisition deficit still remain to be established. Two lines of recent research may be relevant for its explanation: active forgetting mechanisms and memory allocation mechanisms.

Specific cellular and molecular mechanisms of active forgetting were recently subjected to a detailed study in several species (Berry and Davis 2014; Sachser et al. 2016, 2017; Liu et al. 2016; Kitazono et al. 2017). The central idea of the active character of forgetting is the existence of a separate neurobiological mechanism that counterbalances continuous memory acquisition and serves to eliminate unnecessary memory traces (Davis and Zong 2017). Active forgetting has been defined as an intrinsic process that operates constantly in the brain so that new memory consolidation has to overcome it (Davis and Zong 2017). As was found in Drosophila odor conditioning, training triggers both memory acquisition and active forgetting processes. Remarkably, the molecular pathways underlying these two processes are independent and quite different (Shuai et al. 2014). Thus, during pharmacologically induced amnesia both processes can be activated by the training but memory consolidation would be prevented by the drug used, whereas active forgetting mechanisms would be still active. Therefore, administration of a consolidation inhibitor might shift the balance and allow the intrinsic forgetting of specific memory to prevail and prevent the repeated learning.

The second potential explanation of reacquisition deficit relates it to mechanisms of memory allocation. Memory allocation idea refers to an engram being encoded not by a random set of neural elements but rather by specifically selected neurons and synapses (Silva et al. 2009; Frankland and Josselyn 2015). The recruited neurons are selected based on their excitability, which in turn may depend on their age or the level of activated transcriptional factor cyclic adenosine monophosphate–responsive element binding protein (CREB) (Han et al. 2007; Silva et al. 2009; Zhou et al. 2009; Yiu et al. 2014). Increasing the amount of CREB in a subpopulation of amygdala neurons prior to fear conditioning in mice demonstrated an increased probability of the involvement of these neurons in learning-induced neural plasticity (Han et al. 2007). Similarly, in the insular cortex of mice trained for conditioned taste aversion, the subpopulation of neurons with viral-induced CREB overexpression was preferentially recruited into the formation of memory. Accordingly, selective silencing of this subpopulation impaired the retrieval of memory (Sano et al. 2014). Remarkably, the number of amygdala neurons recruited to a given engram remained approximately constant independently of the externally induced changes in the CREB level (Han et al. 2007). These data support the view that neurons are selected to an engram based on a competition process (Han et al. 2007; Silva et al. 2009). Therefore, despite an abundance of potentially available neurons, second training for the same stimulus can be assigned to the same neuronal ensemble that has been engaged in the initial training. If this subpopulation of neurons has been compromised by the drug at the time of training, repeated memory allocation of the specific stimulus might become impossible. Such reacquisition deficit based on selective suppression of memory allocation was reported in mice (Matsuo 2015). Mice that expressed tetanus toxin under the control of *c-fos* promoter were trained in a contextual fear conditioning task. Then, a subset of neurons activated during training was inhibited and memory for the context was tested. Predictably, suppression of the neuronal ensemble resulted in failure of contextual memory retrieval. However, rather unpredictably this inactivation also precluded relearning of the same context, although the animals were able to acquire and retrieve memory for a different context (Matsuo 2015). Based on this data Matsuo (2015) concluded that a neuronal ensemble dedicated to learning is also preferentially involved in relearning and is not replaceable after the initial allocation. In our study, animals were made amnestic during the initial training and the direct influence of the inhibitors had already expired before the second training. However, inability to reacquire the disrupted memory might indicate selective impairment of the neuronal ensemble that was dedicated to the initial learning and is repeatedly recruited in relearning the same stimulus.

We propose to call this phenomenon experience-specific anterograde amnesia, and suggest that it might reflect general properties of memory acquisition, allocation, and impairment mechanisms.

## Materials and Methods

### Animals

Newborn domestic chicks of both sexes (Sussex strain) were obtained from a local supplier (Research and Technological Poultry Institute, Moscow Region) on the day of hatching. They were placed in pairs into metal pens (20 × 25 × 20 cm) and allowed to acclimatize overnight with food and water available. The experimental room was maintained at a temperature of 30°C with a dark/light cycle of 12:12 h. The next morning chicks were pre-trained by two 10-second presentations of a dry metal bead on a rod and only those that pecked at the bead were included in the subsequent experiment (normally >90% of the chicks).

### Behavioral procedures

Chicks of all experimental groups were trained in the one-trial passive avoidance learning task (Lossner and Rose 1983). They were presented with a 2-mm white bead on a rod dipped in a bitter-tasting substance, methyl anthranilate (MeA; Sigma). Typically, after pecking at the bead chicks display a species-specific disgust reaction (head shaking and beak wiping) and subsequently avoid pecking at an identical but dry bead. In all experiments there were four experimental groups. The saline group was trained only once, without administration of any memory blockers; anisomycin and MK-801 groups were trained once along with administration of anisomycin or MK-801, respectively (see below). Two other groups were trained twice: the first training was coupled with a memory-blocker injection, thus producing amnesia, and the second training was given to the amnestic animals either using the same object (the Retraining group) or a novel one, a bead of a different color, red (the Novel group). Time interval between the first and second training was 2 or 24 h, and the retention test for all groups was given in the interval from 0.5 h to 48 h after the second training. For each time point a different group of chicks was used. During the test, chicks were presented for 10 seconds with a dry bead identical to the aversive one used for training (white for the Saline, Anisomycin, MK-801, and Retraining groups; red for the Novel group). Pecking or avoidance of the aversive bead was recorded and a percentage avoidance score was calculated for each experimental group. Finally, the chicks were presented with a dry metal bead that was used in pre-training and served as a neutral (discriminating) stimulus. Only the chicks that pecked at it were included in the analysis. The testing was conducted by an experimenter blind as to which experimental group each chick belonged.

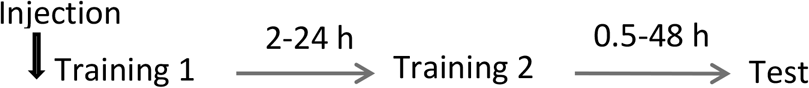

### Injections

An unossified skull of newborn chicks allows the injection of drugs intracerebrally without anesthesia and any obvious changes in the chicks’ behavior (14). The protein-synthesis inhibitor anisomycin (Sigma) was injected intraventricularly using a 10-μl syringe and a headholder designed to direct the injections into the region of the lateral ventricles and adjacent brain areas (Davis et al., 1982). Anisomycin (80 μg/chick, 5 μl/hem) was administered 5 min before the first training. This schedule, as we have previously shown, resulted in amnesia that started after 30 min post-training and lasted for at least 48 h (Anokhin et al. 2002). The non-competitive antagonist of NMDA receptors MK-801 ((+)-MK-801 hydrogen maleate, Sigma) was injected intraperitoneally, 0.4 mg/kg in 0.1 ml saline, or intraventricularly (5 μg/chick, 5 ml/hem) 5 min prior to the first training. Ability of the antagonist to affect recall for passive avoidance in chicks was previously demonstrated both for intraperitoneal and intracerebral administration (Burchuladze and Rose 1992). In all experiments control animals received saline injections. After the experiment, the sites of intracerebral injections were routinely monitored.

### Statistics

Avoidance levels between groups of chicks were compared using the χ^2^ test of independence.

## Acknowledgements

Supported by RFBR#16-04-01848

